# Computational analysis of optogenetic inhibition of CA1 neurons using a data-efficient and interpretable potassium and chloride conducting opsin model

**DOI:** 10.1101/2024.10.17.618665

**Authors:** Laila Weyn, Thomas Tarnaud, Ruben Schoeters, Xavier De Becker, Wout Joseph, Robrecht Raedt, Emmeric Tanghe

## Abstract

Optogenetic inhibition of excitatory populations has been suggested as a novel technique for the treatment of refractory epilepsy. While this approach holds significant potential, achieving seizure suppression in animal models using optogenetics has proven challenging. This difficulty can be attributed to a suboptimal stimulation method that involves numerous complex variables. To provide insight into these parameters, the behavior of the chloride conducting opsin, GtACR2, was fitted to a mathematical description and combined with a conductance-based model of a pyramidal CA1 neuron. The resulting model was adapted to add the ability to study potassium conducting opsins and used to demonstrate that the effect of an optogenetic modulation setup is highly dependent on its parameters and the physiological conditions of the neuronal environment. Stronger inhibition is observed at higher pulse repetition frequencies and duty cycles. Furthermore, potassium conducting opsins were shown to be more stable in use than chloride conducting ones. The dependence of these observations on the opsin model fit was found to be negligible. To determine this, a simplified model (22OMs) is proposed that permits easy implementation of the experimentally derived parameters describing the opsin’s opening and closing dynamics into its mathematical description. This model was also employed to determine that the impact of varying the opsin’s dynamics is insignificant when the opening and closing time constants are altered by a factor between 0.5 and 2. Consequently, this study provides insights into the stimulation and physiological parameters influencing the outcome of an optogenetic inhibition approach and offers a new tool that will facilitate future research into the development of an improved optogenetic modulation protocol for seizure suppression.

## I. Introduction

OPTOGENETICS, a technique that enables control of neuronal activity with light by transducing neurons with gene constructs that encode for light-gated ion channels, ion pumps or receptors, is a very promising neuroscience tool [1]–[3]. The light-induced hyper-or depolarizing currents through these channels can cause inhibition or excitation depending on the opsin type and because the expression of the opsins is controlled by promotors expressed in target neurons, the opsin expression can be cell-type specific [1], [2], [4], [5]. Consequently, whether the modulation will activate or silence neurons can be estimated beforehand, which makes the technique very promising as a treatment option for a variety of neurological disorders, such as drug-resistant epilepsy [2], [3], [6], [7].

The seizures in temporal lobe epilepsy (TLE), one of the most common drug-resistant epilepsies, often originate in the hippocampus. This makes optogenetic inhibition of excitatory hippocampal neurons a possible strategy for TLE seizure suppression [8]–[11] Via optical activation of the halorhodopsin chloride pump, NpHR, expressed in excitatory hippocampal neurons, hippocampal seizure activity was suppressed [10], [12], [13]. However, effective silencing using NpHR requires high opsin expression and continuous illumination at high intensities which can cause unwanted temperature increase [14]–[17] By allowing hyperpolarizing chloride ions through a channel instead of a pump, Guillardia theta Anion-Conducting ChannelRhodopsin 2 (GtACR2) is thought to be a more effective tool [18]. However, lowering the amplitude of epileptic afterdischarges was achieved in vivo, but complete suppression of epileptic activity was not [19]. Furthermore, due to varying chloride ion concentrations between neuronal compartments, GtACR2 can, in certain (epileptic) circumstances, induce excitation instead of inhibition [20], [21]. Newly discovered potassium conducting opsins do not have this problem and have demonstrated their inhibitory power but introducing additional potassium currents may also induce excitation, due to a rise in extracellular potassium levels caused by the opsin currents [14], [15], [22]–[24].

These studies show that the optogenetic approach is promising, but there is definitely room for improvement. Because the chosen opsin influences the waveform of the light induced transmembrane currents and thus the effect of the optogenetic stimulation, this improvement is regularly sought via the discovery and engineering of new opsins with more optimal properties concerning kinetics, biostability, ion conductance and functional wavelength [15], [18], [23], [25]–[30] However, the efficacy of the optogenetic approach depends not only on the opsin but also on physiological properties of the cell, ion homeostasis, the opsin expression level and location, and the illumination properties such as intensity, wavelength and location [31]. These influencing factors require rigorous testing but currently the stimulation paradigm is usually designed using a mixture of trial-and-error and deduction based on data from single cell measurements and knowledge of the mechanisms of action [15], [18]. This is where a computational model can offer a solution, as it enables systematic evaluation of the neuronal response for different opsins under varying circumstances [32].

The mathematical description of an opsin’s behavior, necessary to study its effect in a neuron model, was first published for ChannelRhodopsin2 by Nikolic et al. (2006) [33]. This description can be introduced in conductance-based neuron models that predicate on the Hodgkin-Huxley formalism. Several studies using this approach have been published in recent years, focusing mostly on the understanding and optimization of excitatory applications, not inhibitory ones [32]–[37]. This approach allows the exploration of a vast parameter space and thus can be used to gain insights into the response of opsin expressing neurons to different illumination paradigms allowing for better interpretation and even optimization of the results [32].

In this study, such a combined opsin-neuron model is used to study the effectiveness of an optogenetic inhibitory protocol on a cornu ammonis (CA1) pyramidal cell with a chloride or potassium conducting opsin. This paper offers novel insights into the following aspects of optogenetic inhibition. (i) The impact of the opsin expression level and illumination paradigm is investigated as well as the influence of the intra- and extracellular Cl^−^ and K^+^ concentrations to evaluate the possible occurrence of excitatory effects. (ii) The importance of the opsin model’s accuracy on these simulation results is analyzed to estimate the error caused by uncertainties in the opsin model fit. In order to do this, a simplified version of the opsin model is proposed which is more data-efficient and easily interpretable. (iii) Finally, a sensitivity analysis of the opsin model parameters is performed.

## II. Methods and materials

In this study, a model of a rat CA1 pyramidal neuron is adapted to simulate the neuronal response to optogenetic modulation. The methodology employed to achieve this is described in this section as well as the strategies used for acquisition and analysis of the simulation results.

### A. Opsin model

#### Full opsin model

The photocurrent through the opsin channel is described with the double two-state opsin model (22OM) proposed in Schoeters et al. (2021) [36]. Upon illumination, the opening of the opsin channel is modeled with a potential state transition from closed to open (C → O). During optic modulation, dark-light adaptation causes a change in conductance, which is captured by a transition in states R and S (R → S). When illumination ends, the channel reverts back to its closed, dark-adapted state (O → C and S → R). The state transitions are described by

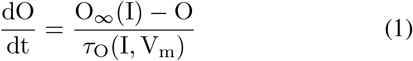

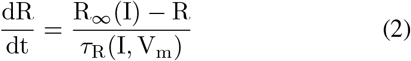

and the state dependent photocurrent of an opsin that is conducting for a specific ion type x is calculated via

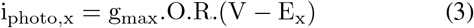

where E_x_ is the reversal potential of the conduction ion type and g_max_ is the maximal specific conductivity for the fully open, dark adapted state. The rectification function G(V_m_), proposed in the original model, is left out because the peak and steady-state currents are approximately linear functions of the holding potential for GtACR2 and additional membrane potential dependency is introduced via calculation of the reversal potential 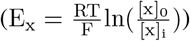 [38].

As illustrated in Figure 1(A), the following features were extracted from experimental data of GtACR2 photocurrents to aid in fitting the model [32]:

**Fig 1:**
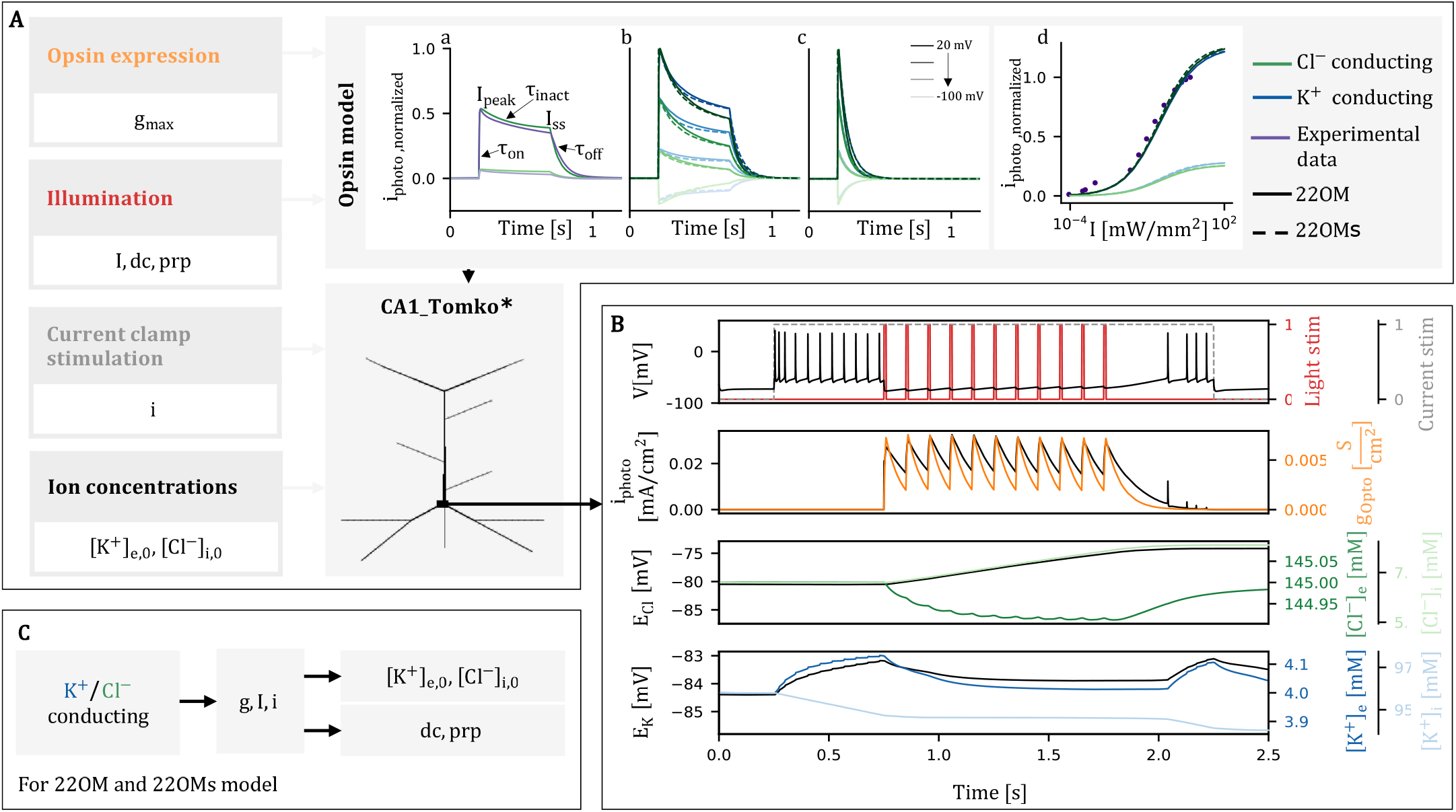
**A**. Model description. Opsin behavior in model and experiments shown under voltage clamp V, initial ion concentrations at rest and g_max_ = 13 10^-3^ mS/cm^2^. All opsin current traces are normalized with respect to peak current of a Cl^−^ conducting opsin for V = 20 mV, I = 4.71 mW*/*mm^2^. (a) V = -20 mV and -60 mV, I = 4.71 mW/mm^2^, t_stim_ = 0.5 s, E_Cl_ constant (b) Varying V, I = 4.71 mW/mm^2^, t_stim_ = 0.5 s (c) I = 4.71 mW/mm^2^, t_stim_ = 10 ms (d) Maximal normalized opsin current as function of light intensity, V = 20 mV and -60 mV, t_stim_ = 0.5 s. **B**. Example of model output for Cl^-^ conducting opsin, i = 0.8 nA, initial ion concentrations at rest, g_max_ = 13 10^-3^ mS/cm^2^, I = 10 mW/mm^2^, dc = 0.1, prp = 100 ms. **C**. Parameter sensitivity study overview.

- Peak current (I_peak_), max current during illumination
- Steady-state current (I_ss_), current during light-adapted state, reached at the end of illumination
- Activation time constant (*τ*_on_), describing current rise from start of illumination until the peak current is reached
- Deactivation time constant (*τ*_off_ ), describing current decay from end of illumination
- Inactivation time constant (*τ*_inact_), describing current decay from I_peak_ to I_ss_
- Recovery time constant (*τ*_recov_), describing decay of I_peak_ in consecutive pulses

The extraction of the time constant features (*τ*_on_, *τ*_off_ , *τ*_recov_, *τ*_inact_) is done using mono-exponential curve fits. The illumination intensity (I) and membrane potential (V_m_) dependency of functions *τ*_O_ and *τ*_R_ are modeled using a log-scale sigmoidal function and logistic regression, respectively [36].

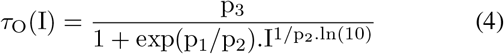

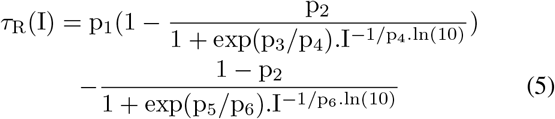

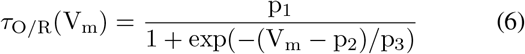

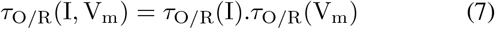

The parameter values p_i_ required to describe these functions are defined separately for each function and derived using a nonlinear least-squares method to fit the extracted time constant features to the following approximations [36].

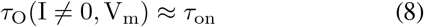

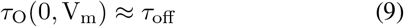

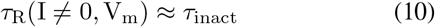

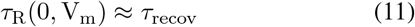

To circumvent restrictions on I_ratio_ imposed by the equation provided by Schoeters et al., *τ*_R_(0, V_m_) was approximated differently. The derivation can be found in the supplementary info.

The equilibrium functions O_∞_ and R_∞_ are only dependent on the illumination intensity and are also fit to a log-scale sigmoidal function defined by 2 and 3 parameters, respectively [36].

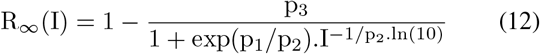

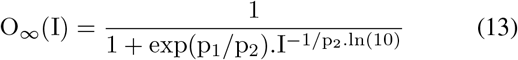

These parameters’ values are determined by minimizing the error between the target values of features I_ss_, I_peak_, 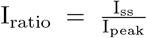 and the model outputs. This is followed by a global bounded particle swarm optimization step during which the error between all feature target values and model outputs is considered. The resulting model comes with 20 fitted parameters, excluding g_max_ and E_ion_.

#### Simplified opsin model

The amount of parameters in the full model make intuitively linking the model parameters to experimental observations quite difficult. Therefore, a simplified version of the double two-state model (22OMs) is proposed in which the approximations mentioned above (eq. (8)-(11)) substitute the complicated *τ*_O_ and *τ*_R_ functions. In the absence of illumination, R_∞_ is 1 and O_∞_ equals 0. When illuminated, R_∞_ is approximated by I_ratio_ because when this is the case, i_photo,x_ of a fully light-adapted open channel is equal to I_ss_ (i_peak_ = g_max_.O_∞_.(R = 1).(V_m_ − E_x_) and therefore i_photo,x_ = g_max_.O_∞_.R_∞_.(V_m_ − E_x_) = I_peak_.I_ratio_ = I_ss_). This results in the following model:

When I = 0

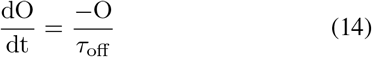

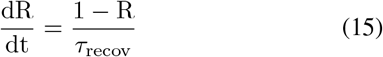

When I ≠ 0

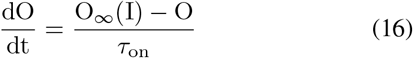

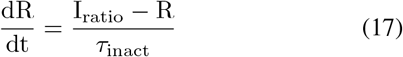

The resulting simplified opsin model no longer has voltage dependent dynamics and the model parameters have been changed to the opsin features I_ratio_, *τ*_on_, *τ*_off_ , *τ*_recov_, *τ*_inact_ and g_max_ in addition to the two remaining parameters defining the log scale sigmoid function describing O_∞_. The default time constants of the 22OMs model are calculated using the formulas of the original model at initial membrane potential V_m,0_ = -65 mV and saturation intensity 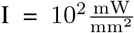 for which the photocurrent plateaus at its maximal value (figure 1Ad). The parameter values can be found in Table I. This simplification approach assumes that the photocurrent shape is independent of V_m_ and I and only scales with I via O_∞_ and V_m_ via the factor (V-E_x_). This model will be used to assess the effect of the opsin model uncertainty on the study results.

#### Data 22OM fit

Single pulse current trace data was acquired from Wietek et al. (2016) [38]. The data is split into two sets. The first, a series of current traces where the holding potential was fixed (V_m_ = 20 mV) and the light flux was varied, is used to determine the intensity dependency of the model, while the other, a series of current traces were the light flux was fixed (I = 4.71 mW*/*mm^2^) and the holding potential was varied (V_m_ = -80 mV through 20 mV in 20 mV intervals) makes it possible to estimate the voltage dependency. Double pulse experiments for estimating *τ*_*recov*_, were acquired from Kopton et al. (2018) [39]. This dataset contains a single data series at a fixed voltage and light flux for GtACR1 as no recovery data was available for GtACR2.

### B. CA1 neuron model

The opsin model is added to each compartment of the reduced-morphology model of a rat CA1 pyramidal neuron described in Tomko et al. (2021) [40] (Fig. 1A). This model was extensively validated with experimental data using HippoUnit, a toolbox that allows for standardized testing of various properties of pyramidal cell models [41]. Consequently, the somatic behaviors are modeled accurately while the reduced complexity results in low computation times.

In order to accurately estimate the effect of the additional optogenetic ion currents on the ion concentrations and vice versa, additional ionic regulation mechanisms are included. The first of these is a Na^+^*/*K^+^ pump, modeled as described in Gentiletti et al. (2022) with 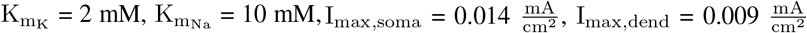 [42].

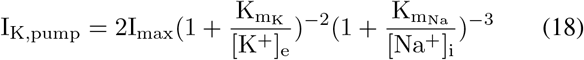

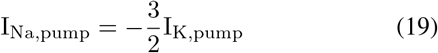

As the major chloride extruder in mature neurons, K-Cl cotransporter KCC2 also plays an important role in maintaining ion homeostasis and is thought to influence epileptic activity [42]–[44] A KCC2 mechanism with strength 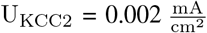 , is implemented via [42]

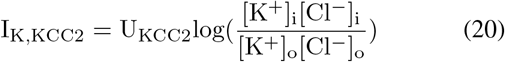

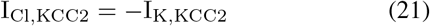

Using NEURON’s rxd module, the neuron compartments are linked to an extracellular matrix with 50 *μ*m grid spacing so the membrane currents impact the intra- and extracellular concentrations of Cl^-^, K^+^ and Na^+^ [45]. As shown in table 1, the resting concentrations can vary in epileptic states. The extracellular concentrations are subject to changes via diffusion.

**TABLE I:**
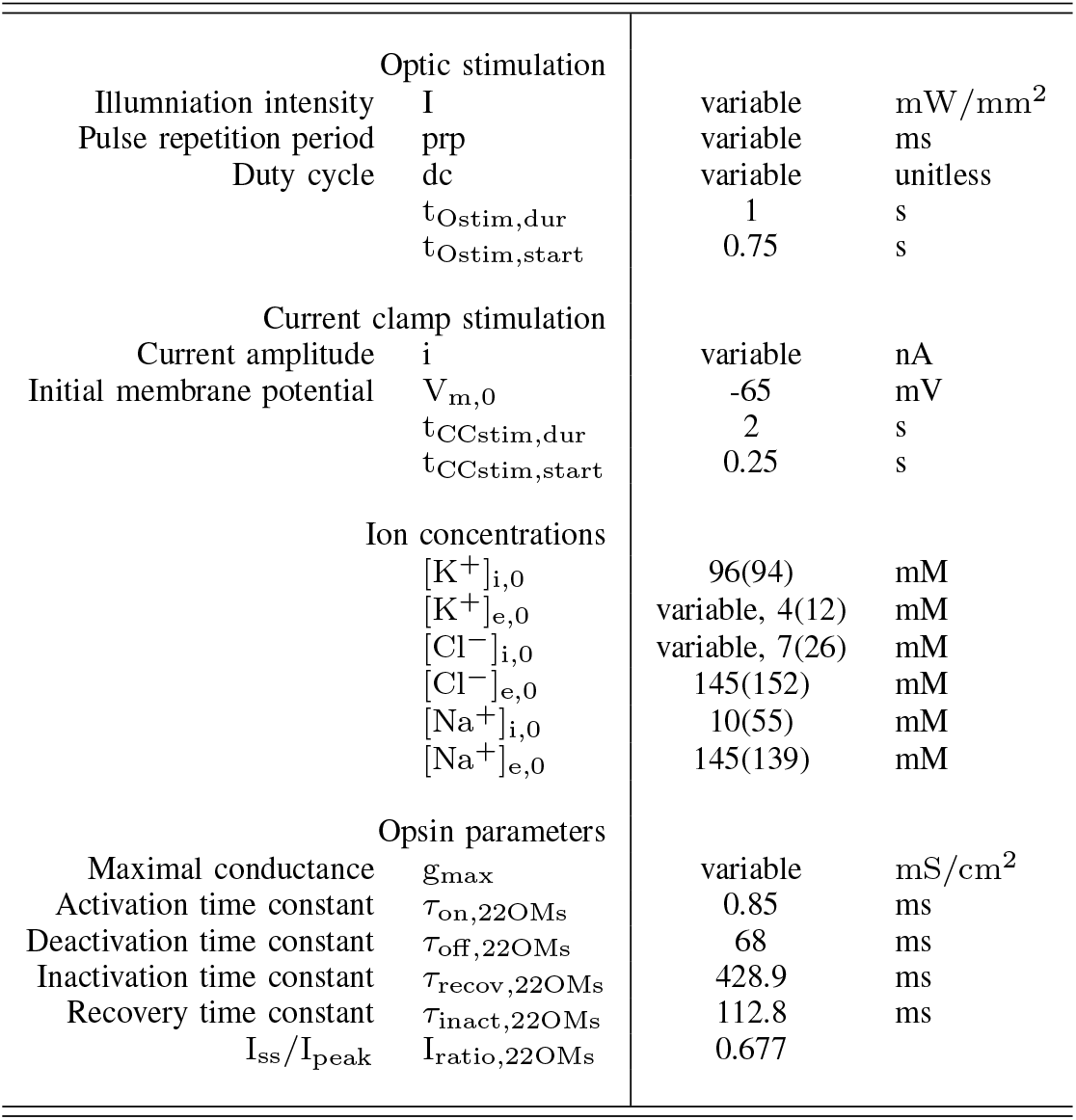
List of important parameters, resting (and epileptic) ion concentrations are given [47].

The diffusion coefficients are: 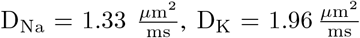 and 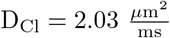 [46].

The behavior of the new CA1 Tomko* model is retested with HippoUnit to evaluate possible changes in performance caused by the alterations. Of the 5 possible tests available in the HippoUnit toolbox (somatic spiking properties, somatic depolarization block, excitatory postsynaptic potential attenuation, action potential backpropagation and synaptic integration at oblique dendrites) solely the two concerning the soma are considered, because only somatic activity is analyzed in this study, meaning the correct simulation of backpropagation and synaptic activity is not relevant [41].

### C. Optogenetic modulation setup

Optogenetic inhibition of the CA1 neuron model is studied by assessing its effect on the neuronal activity resulting from the application of a somatic current clamp of amplitude i. The current clamp starts after 250 ms (t_CCstim,start_) and lasts for 2 s (t_CCstim,dur_). The opsin current is activated by applying a light intensity for 1 s (t_Ostim,dur_), starting 500 ms after the start of neuronal spiking (Fig. 1B). The stimulation amplitude (i), light intensity (I), maximal opsin conductance (g_max_), pulse repetition period (prp), duty cycle (dc) and initial concentrations ([K^+^]_e,0_/[Cl^−^]_i,0_) are variable so their impact on the stimulation’s effectiveness can be determined.

The study described in this work is split into two parts. The first is illustrated in Figure 1C and consists of more than 10^5^ simulations for different parameter configurations, performed for both a K^+^ and a Cl^−^ conducting opsin. The difference in opsin ion type is implemented via the reversal potential used in calculating the photocurrent and by linking this current to changes in the relevant intra- and extracellular concentrations. To limit the simulation time, the study on the impact of the stimulation paradigm is separated from the investigation into the effect of initial ion concentrations: when duty cycle and pulse repetition period are varied, resting initial concentrations are used and varying the initial concentrations is only done for continuous stimulation (dc = 1). Both studies are performed for multiple values of the light intensity I, the maximal opsin conductance g_max_ and the applied current clamp stimulation amplitude i. This decision was made because I and g_max_ directly impact the amplitude of the photocurrent and the stimulation current amplitude is heavily correlated with the amount of photocurrent required for silencing. Therefore, as illustrated in figure 1C, each combination of dc, prp, [K^+^]_e,0_ and [Cl^−^]_i,0_ will be tested for a 3D parameter grid (15Ix9gx8i) with I going logarithmically from 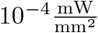 (= photocurrent threshold) to 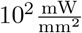 (= photocurrent saturation), g_max_ linearly spanning from 0 to 13 mS*/*cm^2^ (= slightly larger than sodium conductance) and i also ranging linearly from a single spike causing 0.5 nA to an almost depolarization block inducing 1.3 nA.

The second part of this study investigates the effect the opsin model itself has on the results of the previously performed parameter analysis. Therefore, the simulations are repeated with the simplified version of the opsin model (22OMs) and finally an additional parameter study is performed with the parameters (I_ss_, I_ratio_, *τ*_on_, *τ*_off_ , *τ*_recov_, *τ*_inact_) of this model. Initially, the values of I_ss_, I_ratio_, *τ*_on_, *τ*_off_ , *τ*_recov_, *τ*_inact_ are varied separately while the others are kept at the default values listed in table I. The simulations are performed with an injection current i = 0.8 nA and default initial ion concentrations and are repeated for a parameter matrix grid consisting of 3, 2, 2 and 5 values of I, dc, prp and *g*_*max*_ respectively. Finally, the combined effect of *τ*_on_ and *τ*_off_ is studied via simulation with the following parameter settings: 21*τ*_on_ x 21*τ*_off_ x 3I x 3prp parameter grid, i = 0.8 nA, dc = 0.1, g_max_ = 10 mS/cm^2^, default initial ion concentrations.

Simulations were run on a 16GB RAM, quad core processor system. The code was written in python 3.9.12 utilizing NEURON version 8.2.0 [51].

### D. Analysis

The described simulation study setup results in a large amount of data. The analysis methods used to gain insight into the impact of the simulation parameters on the optogenetic modulation results are described in this section.

#### 1) Firing Rate and Normalized Area of Suppression (NAoS)

The effectiveness of the optogenetic inhibition approach is quantified via the firing rate before (FR_init_), during (FR_opto_) and after (FR_post_) illumination. This is defined as the number of spikes that occur during each time interval divided by its duration, each time leaving out a 50 ms transient period. As mentioned in the description of the simulation setup, simulations for each combination of dc, prp, [K^+^]_e,0_ and [Cl^−^]_i,0_ are run for 1080 combinations of I, g_max_ and i. The Normalized Area of Suppression (NAoS) is used to describe the effectiveness of a dc/prp or [K^+^]*/*[Cl^−^] setup with a single value taking into account all different light intensities, stimulation current amplitudes and maximal opsin conductances. As illustrated in Figure 2B, the NAoS is calculated by first determining the minimal g_max_ for which total suppression (FR_opto_ = 0) is achieved for a specific intensity and stimulation current. This results in a line through the intensity-conductance plane. The area to the right of this line is the Area of Suppression. The total NAoS is then calculated by dividing the AoS by the total area of the considered intensity-conductance parameter space and taking the mean over the stimulation current amplitudes.

**Fig 2:**
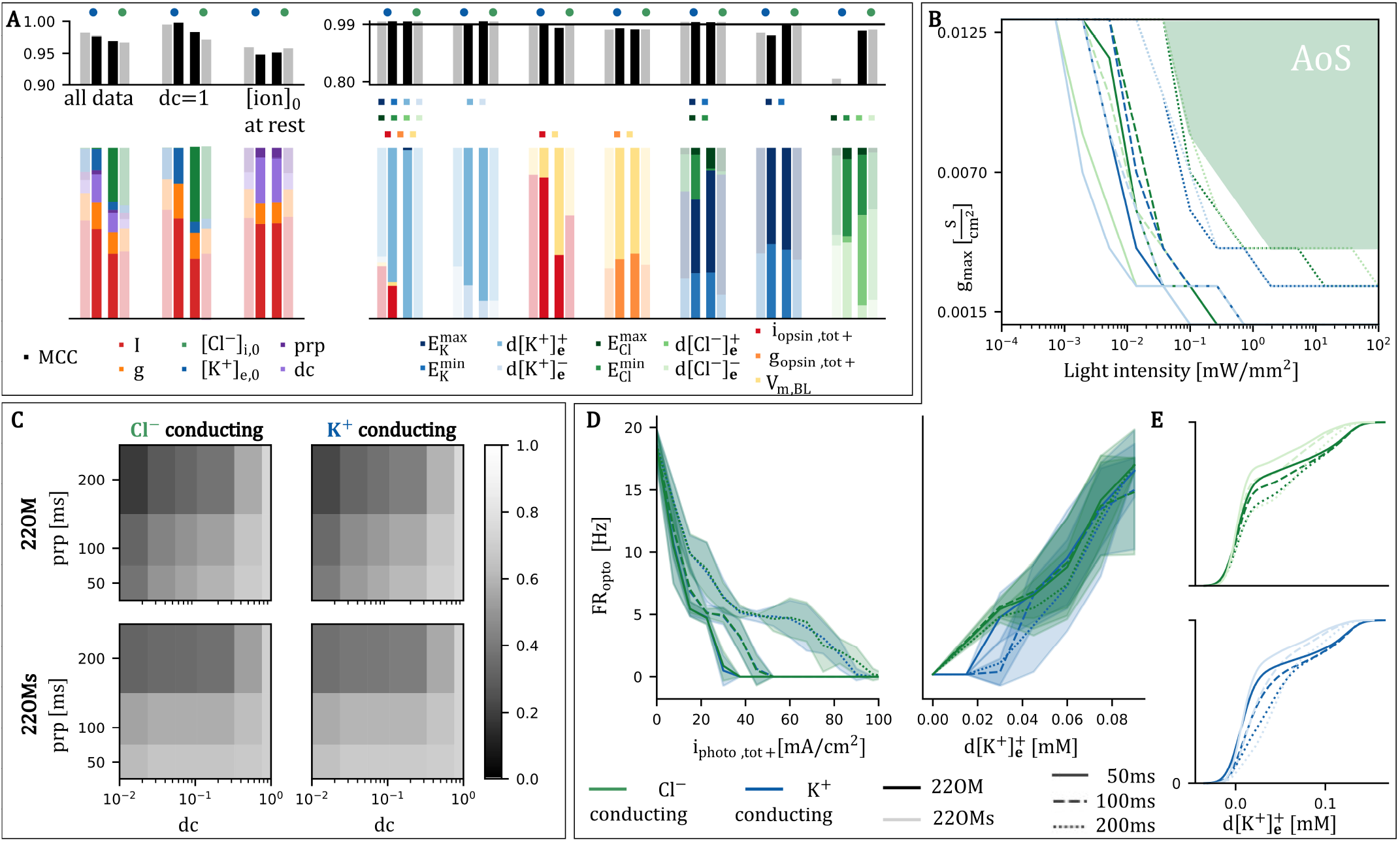
**A**. Performance (top) and PVI (bottom) of classification model for prediction FR = 0 for full/simplified (semi-transparant) 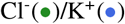 conducting opsin model. (Left) Parameter importance derived from classification model with simulation parameters as features for different data subsets. (Right) Predictive power of physiological metrics as features, varying feature combinations used. Colored squares indicate which metrics are included in each model variant. **B**. Minimal opsin conductance for which the activity resulting from a stimulation current i = 0.8 nA can be completely suppressed as function of the light intensity, dc = 0.14. Area of Suppression (AoS) is illustrated for a Cl^-^ conducting opsin and a pulse repetition period of 200 ms. Initial concentrations at rest. **C**. Normalized Area of Suppression (NAoS) for varying pulse repetition periods, duty cycles and opsin models. Initial concentrations at rest. **D**. Firing rate during illumination as function of the total positive opsin current (left) and total positive change of the extracellular potassium concentration (right) for Cl^-^/K^+^ conducting opsins and different pulse repetition periods. Initial concentrations at rest. Shading indicates standard deviation, over all dc, I, i and g_max_. **E**. Cumulative distribution of total positive change of extracellular potassium concentration during illumination (d[K^+^]_e_^+^). Initial concentrations at rest.

#### 2) Physiological metrics

Aside from the firing rate, which gives information on the effectiveness of the optogenetic inhibition, additional metrics are extracted in order to understand the physiological processes that underlie this neural activity. Because the ion concentrations govern the membrane currents, the total positive and negative change in intra- and extracellular Cl^−^ and K^+^ concentrations 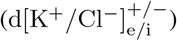 is tracked. The minimum and maximum reversal potential of Cl^−^ and K^+^ that is reached during the illumination period is also recorded 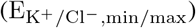, as well as their end values 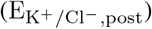.

The direct effect of the optogenetic modulation is quantified by the total positive photocurrent (i_photo,tot+_) and conductance (g_photo,tot_) which is calculated by respectively integrating the photocurrent and opsin conductance, leaving out any negative values. As inhibitory currents decrease the membrane potential, its baseline during AP firing (V_m,BL_) can also serve as a measure of inhibition. It is calculated as the mean of the membrane potential during stimulation, taking only timepoints that are at least 10 ms removed from any action potentials into consideration.

#### 3) Permutation variable importance

A global sensitivity analysis is performed using random forest binary classification as a meta-model [48], [49]. A set of N parameters and/or metrics is given as feature input to the classifier which results in a prediction for a model outcome Y, e.g. FR = 0 or not. The permutation variable importance (PVI) is calculated by randomly shuffling the values of a single feature and measuring the resulting reduction in the model’s predictive performance. This decrease is indicative of the feature’s impact on obtaining the correct prediction so the PVI is used to assess feature importance [48].

The reliability of the PVI values is dependent on the performance of the meta-model which is described using the Matthews correlation coefficient (MCC) [50].

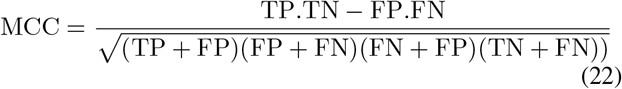

This performance metric combines the results of all four confusion matrix categories (true positives TP, false positives FP, true negatives TN and false negatives FN) in a single value, taking into account possible imbalanced data sets [50]. A random forest classifier consists of 100 trees using the Gini impurity criterion and the final MCC score is the mean of a repeated stratified k-fold cross-validation procedure consisting of 5 folds and 2 repeats.

#### 4) Significance testing

To investigate whether a physiological metric varies significantly between different parameter configurations, a two-side paired Wilcoxon signed-rank test is used. A difference is deemed significant if the p-value is below 0.01.

## III. Results

### A. Opsin model fit

The fitted parameters of the GtACR2 model are summarized in table 2 and a comparison between the experimental and simulated photocurrents is shown in figure 1A. The fitted reversal potential (-66.0 mV) is close to the experimental one (-66.3 mV) [38] but not mentioned in the table because it is replaced in the model by the K^+^ or Cl^−^ Nernst potential. The maximal conductance g_max_ is dependent on the expression levels of the opsin in the cell and is thus treated as a variable.

**TABLE II:** Parameters of opsin model.

| | $\tau_O(I)$ | $\tau_O(V)$ | $\tau_R(I)$ | $\tau_R(V)$ | $O_\infty(I)$ | $R_\infty(I)$ |
| --- | --- | --- | --- | --- | --- | --- |
| p1 | 0.6 | 0.879 | 0.194 | 63.095 | 2.473 | 2.364 |
| p2 | -1.007 | 80.064 | 0.047 | -96.234 | 0.677 | 0.195 |
| p3 | 0.088 | -72.976 | 2.453 | -95.04 |  | 0.323 |
| p4 |  |  | 0.154 |  |  |  |
| p5 |  |  | -0.699 |  |  |  |
| p6 |  |  | 1.522 |  |  |  |

### B. Neuron model validation

The HippoUnit toolbox [41], used to construct and closely match the CA1 Tomko model’s behavior to experimental data, is also utilized to evaluate how the alterations made to enable optogenetic inhibition analysis change the model’s performance. The results are listed in Table III. Lower scores indicate a better biological accuracy and values below 2 are deemed excellent [40], [41]. The modified model, denoted as CA1 Tomko* performs very similarly to the original on the somatic current test but is slightly less accurate in reproducing depolarization block related characteristics. This slight deviation is to be expected as the original model was fitted specifically to perform optimally on these tests. For comparison, the test results for a second experimentally validated but full-morphology model proposed in Migliore et al. (2018) are also added (Table III). The CA1 Tomko* model performs better than this one so the deviations are deemed acceptable.

An additional check was performed to assess the stability of the model without any additional stimulus. After a 250 ms initialization period, the membrane potential changes only 2.5 mV during the rest of the 2.5 s long simulation. The change in E_K_ and E_Cl_ is limited to 0.024 and 0.285 mV respectively and the change of any of the concentrations does not exceed 0.1 mM.

**TABLE III:** HippoUnit scores of original and modified CA1 Tomko model as well as the Migliore full morphology model [52].

|  | depolarization block test | somatic current test |
| --- | --- | --- |
| CA1_Tomko original | 1.72 | 1.07 |
| CA1_Tomko* | 2.29 | 1.02 |
| Migliore | 2.41 | 1.36 |

### C. Parameter analysis

The study of model parameters I, i, g, [K^+^]_e,0_, [Cl^−^]_i,0_, prp and dc is split into 2 parts as shown in Figure 1C. This results in 2 different datasets: one containing data from the concentration study simulation results (dc=1) and another one resulting from simulations performed with [ion]_0_ at rest. For each combination of parameters, this database contains the physiological metrics described in section II.D which will be used to determine the impact of these parameters on optogenetic silencing.

#### 1) Predictive power of parameters and metrics

With the parameters as input features, a random forest classifier predicts whether FR_opto_ is equal to zero or not. The resulting PVI values allow determination of the most impactful parameters. As can be observed in Figure 2A, testing and training with the combined database for which i = 0.8 nA, results in the following importance rankings (≈ if *<* 10% difference): I *>* dc ≈ g_max_ *>* [K^+^]_e,0_ *>* prp *>* [Cl^−^]_i,0_ for a K^+^ conducting opsin and [Cl^−^]_i,0_ *>* I *>* g_max_ ≈ dc *>* [K^+^]_e,0_ *>* prp for a Cl^−^ conducting one. When the classifier data is constrained to a single dataset (either dc = 1 or [ion]_0_ at rest), the impact of g_max_ drops a place in the ranking (except for Cl^−^ conducting dc = 1) but the order importance is otherwise unchanged. The PVI rankings using the 22OMs data are identical to the corresponding 22OM rankings, except for the impact of dc which is consistently lower when the simplified model is used. The model performance is above 0.9 for all models so the resulting rankings are deemed reliable.

The same approach, now with the extracted physiological metrics replacing the input parameters, can be used to assess the predictive power of these metrics. Figure 2A (right) shows that the combination of all metrics contains enough information to almost perfectly (MCC *>* 0.99) classify all data. The total positive change in extracellular potassium 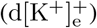 has the highest predictive power and even when the total positive and negative change in extracellular potassium are the only two metrics used a performance score above 0.99 is reached for all opsin models. Intuitively and according to the PVI values of the classifier using all metrics, high importance for V_m,BL_ and i_opsin,tot+_ is expected. This is confirmed by the above 0.99 performance value for 3 of the 4 opsin models when only these metrics are used. The model’s predictive ability drops when the conductivity of the opsin is used instead of the current. Using the reversal potential metrics also results in a well performing classifier (MCC *>* 0.99). The drop in performance when the chloride reversal potentials are left out shows that they hold valuable information but potassium based metrics clearly hold more predictive power as is shown by the low performance of the classifier containing only chloride metrics, especially for K^+^ conducting opsins.

#### 2) Optic stimulation parameters

The importance ranking indicates that the pulse repetition period has a low impact on the modulation result. However, figure 2B shows that there is an impact nonetheless. For a constant dc = 0.14, a simulation with longer off-time between pulses requires a higher light intensity to achieve complete silencing with the same opsin conductance. This trend is also visible in figure 2C, which indicates that lower prp values result in more effective silencing for all opsin models and dc values (dc ≠ 1). Furthermore, this figure confirms that an increase in dc improves the optogenetic silencing. The same trends are observed when inspecting the 22OMs model, but the influence of decreasing dc is less pronounced. The total sum of AoS over all i/dc/prp combinations is 80 (78), 84 (82) for the (simplified) Cl^−^ and K^+^ conducting opsin model respectively, indicating a slightly better overall performance of the latter.

Intuitively, the positive impact of a decreased pulse repetition period could be explained by the fact that this coincides with more pulses and thus a higher total photocurrent due to slow closing induced photocurrent tails. A Wilcoxon rank test does indeed indicate a higher mean photocurrent for faster pulse rates (p*<*0.01 for all pairs) and Figure 2D shows that there is a correlation between FR_opto_ and the total photocurrent. However, this figure also shows that a similar current results in higher firing rates when prp increases, indicating that the increased photocurrent hypothesis is insufficient. Alternatively, figure 2D shows that there is a correlation between larger positive changes in [K^+^]_e_ and FR_opto_ without a visible effect of prp. As can be seen in figure 2E, the 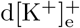 values are significantly higher for lower pulse frequencies (p*<*0.01 for all), indicating that 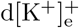 could be an explanatory metric.

#### 3) Ion concentrations

Figure 3A shows how varying ion concentrations influence the effectiveness of optogenetic silencing. The potency of Cl^−^ conducting opsins depends heavily on both the extracellular K^+^ and intracellular Cl^−^ concentration. High values of either can result in failure of silencing and, as illustrated in figure 3B, a raised baseline potential (V_m,BL_), indicating increased excitability, can occur when [Cl^−^]_i,0_ is elevated and [K^+^]_e,0_ is not. Unlike the Cl^−^ conducting opsin model, the K^+^ depending one is not affected by a varying [Cl^−^]_i,0_ concentration but the effect of a [K^+^]_e,0_ increase is the same. From Figure 3B can be inferred that a potassium photocurrent always has an inhibitory effect. A bigger drop in baseline potential is observed for higher initial values of [K^+^]_e_ which can be attributed to this increased concentration elevating the initial baseline potential and the photocurrent returning it back to normal.

**Fig 3:**
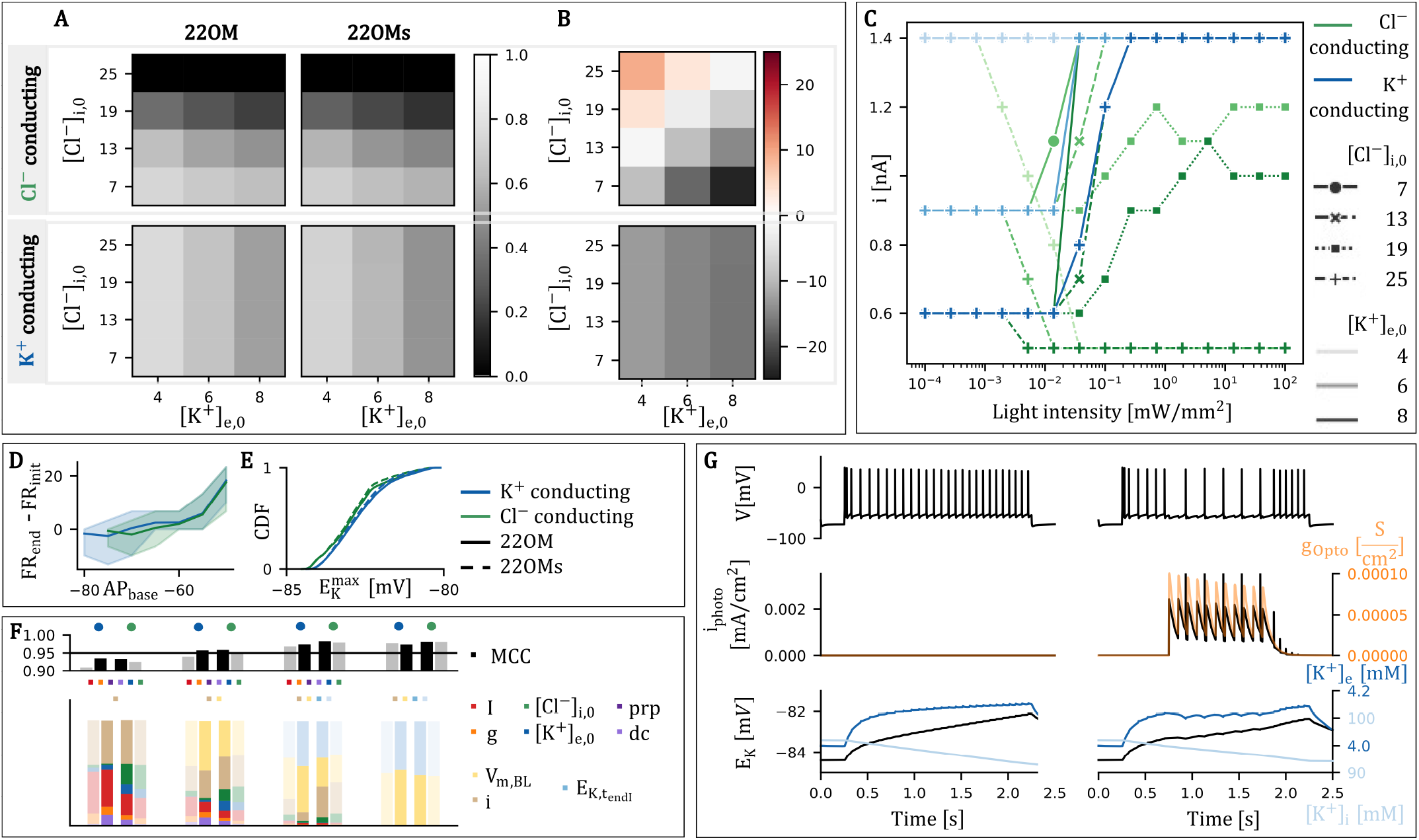
**A**. Normalized Area of Suppression for varying initial Cl^-^/K^+^ concentrations and opsin models (dc = 1). **B**. Mean difference in baseline potential 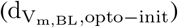 during and before illumination in mV. **C**. Lowest current stimulation causing depolarization block for any opsin conductance, as function of I (dc = 1). **D**. Difference in firing rate before and after illumination as a metric of hyperexcitation, as function of V_m,BL_ during illumination. Shading indicates standard deviation. **E**. Cumulative distribution of maximal K^+^ reversal potential reached during illumination. **F**. Performance (top) and PVI (bottom) of classification model for prediction of FR_end_ -FR_init_ for full/simplified (semi-transparant) 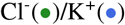 conducting opsin model, varying feature combinations used. Results in which a depolarization block occurs at any point were not included. Colored squares indicate which metrics are included in each model variant. **G**. Effect of K^+^ conducting opsin on a 0.75 nA stimulation current, with (right) and without (left) illumination. prp = 100 ms, dc = 0.1, initial concentrations at rest, g_max_ = 10^-3^ mS/cm^2^, I = 0.01 mW/mm^2^

The elevation of the initial baseline potential can cause the neuron to lose its ability to fire and thus enter depolarization block (DB if AP_BL_ *>* -45 mV and FR = 0 Hz). The effect of optogenetic stimulation on a neuron that has entered DB is displayed in figure 3C. For [K^+^]_e,0_ = 4 mM, DB occurs only when the light intensity and [Cl^−^]_i,0_ are sufficiently high to induce an excitatory Cl^−^ photocurrent ([Cl^−^]_i,0_ ≥ 25 mM and *I >* 10^−1^). For higher [K^+^]_e,0_, initial DB occurs when i reaches a threshold value (0.9nA and 0.6nA for [K^+^]_e,0_ = 6 mM and 8 mM, respectively). In these cases, stimulation at sufficient intensity of a K^+^ conducting opsin can restore the neuron to its resting membrane potential regardless of the current injection. A Cl^−^ conducting opsin is only able to do the same when the intracellular chloride concentration is low.

### D. Increased excitability

The previous paragraph reported that, unlike increased [Cl^−^]_i,0_ in combination with a Cl^−^ conducting opsin, elevated extracellular potassium levels do not impact the effectiveness of K^+^ photocurrent induced silencing and increased excitability during illumination is not observed. However, a K^+^ photocurrent affects the potassium concentration which means elevated [K^+^]_e_ induced hyperexcitability is a possibility. Figure 3D shows that hyperexcitation in the form of an increased firing rate after stimulation in relation to its initial value can occur (dFR_init,post_), specifically when the silencing protocol is less effective as indicated by a high baseline potential. This indicates that failure to inhibit might result in hyperexcitation due to elevated [K^+^]_e_ caused by a potassium photocurrent. However, this is not the case because the difference in firing rate for high baseline potentials also occurs when the photocurrent is caused by Cl^−^ ions and the distributions in figure 3E show E_K_ reaches the same maximal value for both Cl^−^ and K^+^ conducing opsins.

A new random forest classifier with dFR_init,post_ equal to or differing from zero as outcome variable is trained and tested with different parameters and metrics to gain more insight into the occurrence of hyperexcitation. Simulations during which a DB was established at any point were not included. Using only input parameters i, I, g_max_, [K^+^]_e,0_, [Cl^−^]_i,0_, prp, and dc is not sufficient to acquire a model with above 0.95 MCC. From Figure 3F, it can be deduced that i and I hold the most predictive power in this model which is in line with the observation that hyperexcitation occurs when suppression fails. Adding V_m,BL_ as a feature increases the model’s MCC but only when the end of stimulation potassium reversal potential is introduced 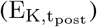, does the MCC value reach 0.95 for each opsin model. Using only V_m,BL_ and 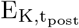 results in an even better model (MCC*>*0.97). This suggests that the hyperexcitation is due to a rise in E_K_ and more likely to occur when the inhibition is less effective (thus causing a higher V_m,BL_), but does not give insight into why E_K_ rises for both K^+^ and Cl^−^ conducting opsin channel. Therefore, the answer has to be found in analysis of the evolution of the potassium concentration over time during illumination (Fig. 3G). These graphs show that E_K_ rises due to the neuron’s activity and the firing rate increases even when there is no optogenetic stimulation. Furthermore, E_K_ drops slightly during the illumination pulses, indicating an inhibitory effect but restores during the inter-stimulation periods. After the illumination, E_K_ continues its previous upward trajectory, increasing the excitability and causing the same heightened firing rate that would have been achieved without illumination. Lastly, it is important to note that the maximal amount of potassium extruded due to a photocurrent in this study (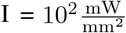 , g = 13 mS/cm^2^, dc = 1) is only half the amount that is extruded by regular, naturally occurring channels with the same settings but without the illumination.

### E. Sensitivity opsin model

As mentioned in the previous sections, the observations on parameter influence do not vary when the opsin model is replaced by its simplified version which indicates that the outcome of the optogenetic stimulation is not overly sensitive to small changes of the opsin model. The impact of opsin parameters I_ratio_, *τ*_on_, *τ*_off_ , *τ*_recov_ and *τ*_inact_ is studied using this simplified model. The results of an experiment where the values of these parameters were varied separately are illustrated in figure 4A. They indicate that none of these parameters have a significant influence when illumination is continuous and only changes to *τ*_on_ and *τ*_off_ cause a larger than 2.5 Hz mean deviation of the resulting FR_opto_. When *τ*_off_ is too small, the stimulation can lose its inhibitory effect entirely because the short pulses are not enough to suppress the activity when the photocurrent ceases completely in between. A deviation of *τ*_on_ induces significant changes once its value exceeds the illumination pulse duration (10-100 ms depending on prp and dc) and the channel does not have the time to open completely, resulting in a lower photocurrent. From figure 4B, it can be concluded that even though altering *τ*_on_ and *τ*_off_ can impact the results significantly, the observations concerning the effect of prp, dc, I and g_max_ (i.e., superior silencing can be achieved by increasing dc, I, g_max_ and decreasing prp) remain valid through these alterations.

**Fig 4:**
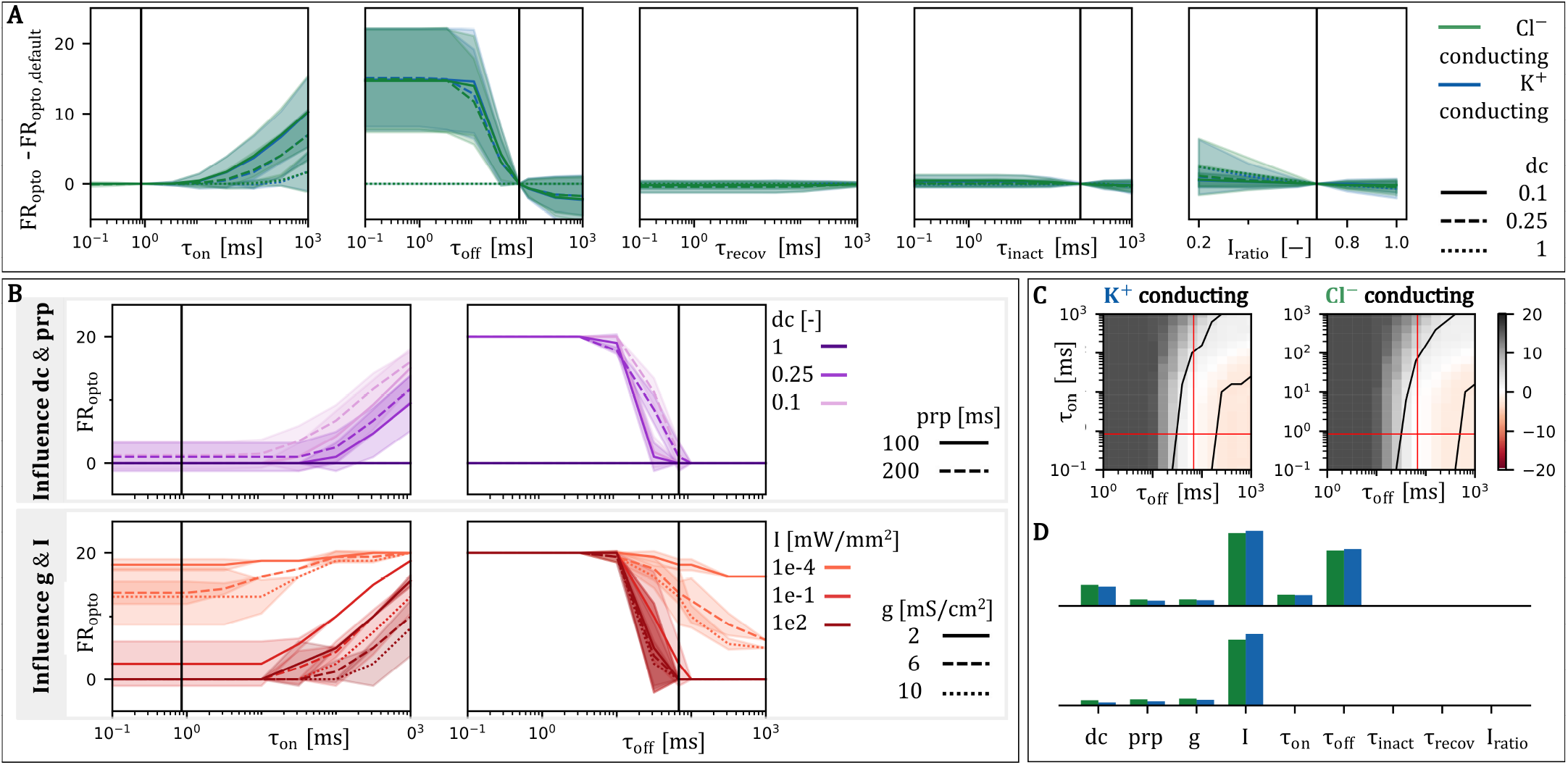
**A**. Change in firing rate during stimulation induced by changing opsin model parameters separately. Shading indicates standard deviation of results from varying I, g_max_ and prp. (i = 0.8 nA, default initial ion concentrations) **B**. Firing rate during illumination as a function of *τ*_on_ and *τ*_off_ for different I, g, dc, prp. Shading indicates standard deviation over results with varied values of not separated variables (i = 0.8 nA, default initial ion concentrations). **C**. Mean difference in firing rate (in Hz) w.r.t. default parameters, averaged over varying I and prp for different combinations of *τ*_on_ and *τ*_off_ . Red lines: default values. Between black lines: dFR *>* 2.5Hz (i = 0.8 nA, dc=0.1, g_max_ = 10e-3 mS/cm^2^, default initial ion concentrations). **D**. PVI of classification models for prediction FR = 0 using simulation and opsin model parameters as features for 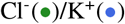 conducting opsin model. (upper) all *τ*_on_ and *τ*_off_ values used. (lower) only *τ*_on_ and *τ*_off_ bigger than half and smaller than double their default value taken into account. MCC *>* 0.99 for both models.

The combined effect of *τ*_on_ and *τ*_off_ on the model results is illustrated in figure 4C, indicating that an alteration with factor 0.5-2 in both parameters is allowed before the deviation of the firing rate exceeds 2.5 Hz. A random forest classifier with all varying parameters as features and FR_opto_ = 0 or not as output is once again used to estimate their importance. The results in figure 4D indicate that I, dc, *τ*_on_ and *τ*_off_ play the biggest part with the illumination intensity and opsin closing time more important than the others. This indicates that uncertainty in *τ*_off_ and, to a lesser extent, *τ*_on_ can impact the results significantly. However, when the possible *τ*_on_ and *τ*_off_ values are constrained to 0.5-2 times the default value, they lose their predictive value and the illumination intensity becomes the most important once again, followed by dc, g_max_ and prp.

## IV. Discussion

In this work, simulation studies were performed to gain insight into the effect of various physiological and optical parameters on the efficacy of optogenetic inhibition of a CA1 pyramidal neuron. Understanding the impact of these parameters can aid in the development of more efficient strategies for optogenetic suppression of seizures in the treatment of TLE and unveil possible disrupting factors. Furthermore, the proposed simplified opsin model can provide insight into desirable opsin characteristics and may facilitate *in silico* experiments with many different opsins in other applications.

### A. Modeling opsin characteristics

It is assumed that accurately modeling the effect of an optogenetic stimulation setup requires a precise mathematical description of the opsin’s behavior. In this study, the opsin model was constructed by fitting the 22OM model on experimental data obtained from the chloride conducting opsin GtACR2 [18], [36]. Due to limited data availability this could not be done optimally. There was only a single set of photocurrent traces for different membrane holding potentials and another set for varying light intensities, meaning that possible cross dependencies were not taken into consideration [38]. Furthermore, the single current trace used to determine *τ*_recov_ came from the slightly different opsin GtACR1 and the rectification function G(V_m_) included in the original model described in Schoeters et al. (2021) was set to one [36], [39]. These limitations could theoretically impact the accuracy of the model. However, the fitting procedure was deemed acceptable because a good match to the available data was achieved (see figure 1A) and this study focuses not on GtACR2 specifically but on the impact of multiple parameters on a generic K^+^ or Cl^−^ conducting opsin.

The newly proposed simplified opsin model (22OMs) is used to determine whether the insights gained via simulations depend heavily on the model fit. The observations concerning the influence of I, g_max_, prp, dc, [K^+^]_e_ and [Cl^−^]_i_ discussed in the previous section remain unchanged when the original opsin model is exchanged for the simplified one. The only remarkable difference lies in the impact of decreasing dc which is lower for the 22OMs model (figure 2C). Additionally, incorrect estimation of *τ*_recov_, *τ*_inact_ and I_ratio_ were shown to have little effect on the optogenetic stimulation outcome. However, the impact of a simultaneous variation of all parameters was not considered in this study, and it is expected that the influence of *τ*_recov_ and *τ*_inact_ will increase at lower I_ratio_. The limited impact when the deviation factor of *τ*_on_ and *τ*_off_ lies between 0.5 and 2 shows that there is a significant margin of error allowed in their estimation as well. The design of the 22OMs model assumes that the time constants are independent of V and I. When looking at the extracted data features *τ*_on_ and *τ*_off_ for different voltages, this appears to be the case. For decreasing intensity levels, *τ*_off_ also decreases but its values remain within the allowed range. *τ*_on_ however becomes more than a factor 2 larger when low intensities are applied. This implies that the 22OMs model loses some reliability when low illumination intensities are used. It also explains why decreasing dc has more impact when the 22OM model is applied because the higher *τ*_on_ values that occur with this model cause lower photocurrents to be reached during a short illumination time.

These observations indicate that the simplified model can facilitate future research into specific opsins because its application results in minimal deviations of the simulation outcome. Additionally, it can be fitted using opsin data that is conventionally reported when presenting a new opsin: a single photocurrent trace at high, close to threshold, illumination intensity, a multiple pulse experiment and the i_photo_ vs I curve. The model contains less differential equations as well as a lower amount of easily extractable parameters than other previously reported models [33], [35], [36], [57], [58]. I_ratio_ can be directly read from the photocurrent trace and time constants *τ*_inact_, *τ*_on_ and *τ*_off_ can be fitted to it using mono-exponential curve fits. *τ*_recov_ can be mono-exponentially fit to the multiple pulse current trace. This experiment is not always performed but as shown in this study, deviation of *τ*_*recov*_ does not significantly influence the results. Parameters p_1_ and p_2_ can be extracted by fitting the O_∞_ function on the normalized i_photo_ vs I curve. Extraction of g_max_ is the hardest as it is a combination of the single channel conductance and the expression level [37], [56]. It can be done by matching the model’s calculated photocurrent to the I_peak_ value but this is not a straightforward procedure because the model uses units per cm^2^ for both g_max_ and i_photo_ while I_peak_ is usually reported in nA for a cell with unreported area. Furthermore, the expression level can vary significantly between cells and is rarely quantified. The only reported value, extracted from bacteriorhodopsin, is 1.3e10 channels/cm^2^ but indirect estimation has resulted in 4.4e12 channels/cm^2^ [59], [60]. Moreover, reports for the much better investigated ChannelRhodopsin2 (ChR2) suggest that the single channel conductance can vary between 40 and 250 fS [37], [56], [58], [61]–[63]. This data implies that g_max_ may vary between 0.52 and 1100 mS/cm^2^ for a single opsin type, indicating a need for better quantification efforts before *in silico* models can be used for the exact optimization of optogenetic stimulation applications.

In conclusion, the 22OMs model makes it easier to design *in silico* experiments with (new) specific opsins but one must always be aware of its limitations. Firstly, g_max_ may be estimated but must be treated as unknown. Secondly, most accurate performance will be achieved for high light intensities. Furthermore, the modeled relationship between the intensity and the photocurrent is not voltage dependent. In fact, the only V dependence is introduced via the (V_m_ − E_x_) factor. While there is no consensus on the voltage dependence of the opening-closing dynamics, it cannot be ruled out and thus must be kept in mind. All time constants are independent of I and V_m_ which can cause inaccuracies in the simulation of short, low-intensity pulses. Besides these limitations, this model has the upside of avoiding a complex, difficult to understand overfit on limited data.

### B. Optimization of optogenetic silencing

The simulation results clearly indicate that each of the investigated parameters can influence the effectiveness of an optogenetic inhibition paradigm, meaning that all these parameters must be taken into consideration when optimizing the stimulation setup.

The illumination intensity I has the most significant impact on the effectiveness of the optogenetic inhibition paradigm. Continuous stimulation at saturation intensity is guaranteed to result in the strongest suppression. This type of stimulation is however not advisable because tissue illumination causes heating. A single 100 ms pulse at saturation intensity 100 mW*/*mm^2^ can already induce a temperature increase of 0.5 °C and a 2 ^*°*^C rise can induce changes in neural parameters and thus brain activity [37], [53]–[55]. Using pulsed stimulation decreases power input and thus the temperature increase but lowering the duty cyle also decreases the suppression effect.

Lowering the pulse repetition period allows for better inhibition while maintaining the same power. Results indicate that this effect cannot be attributed to additional currents due to slow closing induced photocurrent tails. A study on the predictive qualities of various metrics suggests that the total amount of extruded potassium is an explanatory metric. This is likely due to the link between this outward flow of K^+^ and the membrane potential, making the extrusion of K^+^ a predictor for the level of inhibition. The stimulating current clamp raises the membrane potential (i = dV_m_*/*dt), inducing an increased current i_K_ ∼ (E_K_ − V_m_) and thus causing more extrusion of K^+^. Similarly, inhibition coincides with a decrease of the membrane potential and thus a lowering of i_K_, giving [*K*^+^]_*e*_ the time to restore and decrease. (This behavior can be observed in figure 3G.) As mentioned in section III.D, the photocurrents of a potassium conducting opsin are small compared to the other, naturally occurring potassium currents so their effect is negligible. Therefore, the lower amount of change in [K^+^]_e_ when the prp is decreased can be interpreted as the membrane potential not rising fast enough between pulses to reach the firing threshold. This supports the hypothesis that a minimum membrane potential — and therefore maximal inhibition — is reached during illumination, independent of pulse duration. Consequently, whether firing occurs between pulses depends on the time available for the neuron’s membrane potential to recover, which is determined by pulse repetition period.

Theoretically, an optimal stimulation paradigm (I/dc/prp) to maximize optogenetic suppression using a specific opsin, while minimizing temperature increase and power delivery, can be found using *in silico* experiments and optimization algorithms. However, results indicate that the level of suppression depends on the maximal opsin conductance so it can be inferred that the optimal combination of I, dc and prp in an optogenetic setup may too. A high g_max_ value is preferable for effective inhibition but to find optimal values of the other parameters g_max_ must be known. As explained in the previous section, this is very challenging, which limits the application possibilities of the model.

Choosing the right opsin is also part of the optimization process. Results indicate that a potassium conducting opsin is slightly more effective in silencing the CA1 pyramidal neuron than a chloride conducting one. This can be explained by E_K_ being lower than E_Cl_ at rest, resulting in a higher total photocurrent for the same set of parameters which is confirmed by a photocurrent distribution analysis and Wilcoxon rank test (p*<*0.01) [47]. Furthermore, simulation results support the hypothesis that a chloride conducting opsin can induce hyperexcitation when distorted ion concentrations occur, leading to the conclusion that a K^+^ conducting opsin is the better, more stable choice [15], [20]. As seizure-induced distorted Cl^−^ concentrations can occur in all subcellular compartments, soma-targeting of the opsin is not a viable solution for TLE treatment applications [20]. The on and off dynamics of the opsin also play an important role. In an ideal optogenetic stimulation setup *τ*_on_ and *τ*_off_ are respectively much smaller and longer than the illumination pulse duration. The observed positive effect of increasing *τ*_off_ is quite limited (only 4Hz for a factor 1000 increase) but this effect may vary when studied for a larger set of pulse frequencies.

### C. Risk of hyperexcitability

As inhibition is the goal, hyperexcitation should definitely be avoided. Theoretically, potassium photocurrents could however lead to an increase in extracellular K^+^ levels and E_K_, resulting in higher excitability. The simulation results show that the total K^+^ photocurrent is always smaller than what would be extruded without optogenetic intervention and E_K_ consistently decreases during illumination. Any hyperexcitation, linked to E_K_ increase, that occurred in the simulations can be attributed to the continuously applied current clamp stimulation in combination with incomplete suppression. A rise in [K^+^]_e_ and thus E_K_ is then caused by the increase in membrane potential due to the excitatory current input. However, these results do not take into account the fact that recovery of extracellular potassium levels is more challenging in a network where many neurons contribute to the build-up nor the possibly reduced efficiency of potassium buffering in epilepsy. Therefore, we can conclude that it is unlikely for photocurrents to induce hyperexcitation through changes of ion homeostasis but the effect in an epileptic hippocampal network requires further investigation.

Hyperexcitation can also occur when, by distorted initial ion concentration, altered reversal potentials impact the photocurrents. High intracellular Cl^−^ concentrations result in hyperexcitation due to E_Cl_ exceeding the resting membrane potential leading to an outward, excitatory photocurrent (i_photo_ ∼ (E_Cl_ − V_m_)). The same effect does not occur for K^+^ conducting opsins even though E_K_ does rise above the membrane’s resting potential when [K^+^]_e,0_ is increased. This can be explained by the increase of [K^+^]_e,0_ also causing a simultaneous rise in V_m_ meaning (E_K_ −V_m_) retains the same sign so the photocurrent remains inhibitory.

### D. Limitations and future work

The computational model designed for this study was used to gain useful insights into the influence of stimulation parameters and physiological circumstances on optogenetic suppression. As in all computational replicas of natural phenomena, some approximations were made that should be taken into consideration when interpreting the result. Firstly, while the results of this work indicate that the 22OMs model provides a good, straightforward approach to accurately model the behavior of opsin dynamics based on easily gathered, small amount of data, its versatility and general application abilities should be further investigated. Also, due to its aforementioned limitations, this opsin model cannot be used to identify the exact optimal settings of an optogenetic setup.

Secondly, while the CA1 Tomko* model was validated on experimental data, it remains an approximation of the truth and does not take into account any inter cell variability caused by variations in complex morphology and channel conductances, the influence of which should be further investigated [52]. On the other hand, less complex, one-or two-compartmental models of CA1 pyramidal cells also exist and require less computational power [64], [65]. Since temporal lobe epilepsy is a brain network disorder, optogenetic suppression of TLE seizures is more complex than simply inhibiting a single CA1 pyramidal neuron. Therefore, further investigation into optogenetic inhibition of hippocampal network activity is required. For the design of such an *in silico* network study, it would be useful to know the impact of replacing the CA1 Tomko* model with one of these simpler neuron models on the model outcome.

Finally, the illumination devices typically used for optogenetic applications are quite small and light propagation in neural tissue is not homogeneous. Therefore, the assumption of uniform illumination intensity at every neuronal compartment is another gross approximation [37], [53]. Because this study focuses on somatic behaviors, this was not deemed an issue. Important to note is also that the illumination intensities reported in this work are the ones at the neuron itself meaning higher intensities should be applied at the device in order to get the same effect at the neuron location. In any future network investigations, the spatiotemporal dynamics of the optic field should be taken into account via light and heat propagation modeling [16], [37], [66]. Using this approach, light intensities can be calculated at each point relative to the source location and the local temperature increase can be calculated. This way the optimal illumination profile and location(s) can be investigated. The applicability of this approach is however hindered by uncertainty of the tissue’s optical properties [37], [67], [68].

## V. Conclusion

In this study, a computational model of a CA1 pyramidal neuron was adapted to include optogenetic functionality and used to study the impact of various parameters on the effectiveness of an optogenetic inhibition paradigm. The insights gained into the influence of stimulation parameters illumination intensity, pulse frequency and duty cycle were in line with expectation. Longer, more intense pulses at a high frequency proved most effective for suppressing neuronal activity. Whether total inhibition is achieved is also largely dependent on the applied current clamp as well as the opsin conductance and expression level. Consequently, precise measurement of single channel opsin conductance is necessary before the model can be employed to find a solution for optimal silencing at lowest power thus minimizing temperature increase. In comparison to chloride conducting opsins, potassium conducting ones are shown to be slightly more effective and more robust to seizure induced changes in ion concentrations. Furthermore, no evidence was found of potassium photocurrent induced hyperexcitability.

The uncertainty introduced to the model via imprecise fitting of the opsin behavior does not influence the results described above which remain unchanged for a wide range of the opsin model’s parameters. Of those parameters, only the opening and closing time have a significant effect on the suppression. Optimal photocurrents are achieved when *τ*_on_ and *τ*_off_ are respectively much shorter and longer than the illumination pulse duration. Furthermore, the simplified model allows direct implementation of a different opsin’s behavior without complex fitting procedure, instead using the experimentally accessible time constants, the maximal conductance and relationship between the illumination intensity and normalized photocurrent. This data-efficient approach makes the model easily interpretable and adaptable so it can facilitate future research concerning the determination of the best opsin for a specific application or aid in the optimization of a stimulation setup for a chosen opsin in a single neuron or network model.

## VII. Acknowledgment

We would like to thank Dr.Jonas Wietek and Dr.Franziska Schneider-Warme for sharing their experimental data on GtACR2.

