## Supplementary Info for "Computational analysis of optogenetic inhibition of CA1 neurons using a data-efficient and interpretable potassium and chloride conducting opsin model"

### I. Derivation of $\tau_{\mathrm{R}}(\mathrm{I}=0)$

In Schoeters et al. (2021), the time constant $\tau_{\mathrm{recov}}$ that describes the recovery of $\mathrm{I_{peak}}$ over multiple pulses is defined by the following equation:

$$\mathrm{I(t)} = \mathrm{I_{peak_1}}[1-\exp(-\frac{\mathrm{t}-\mathrm{t_{off_1}}}{\tau_{\mathrm{recov}}})] \tag{1}$$

Here the first peak ($\mathrm{I_{peak_1}}$) is maximal and a second peak ($\mathrm{I_{peak_2}}$) can be defined so that $\mathrm{t_{peak_2}}$ - $\mathrm{t_{off_1}}$ = $\tau_{\mathrm{recov}}$ and thus $\mathrm{I_{peak_2}}$ = $\mathrm{I_{peak,1}}(1-\mathrm{e}^{-1})$. This approach implies that peak values lower than steady state occur when

$$\mathrm{I_{ss}} > \mathrm{I_{peak_1}}[1-\exp(-\frac{\mathrm{t}-\mathrm{t_{off_1}}}{\tau_{\mathrm{recov}}})] \tag{2}$$

$$\Leftrightarrow \mathrm{t} < \mathrm{t_{off_1}} + \tau_{\mathrm{recov}}\ln(\frac{1}{1-\mathrm{I_{ratio}}}) \tag{3}$$

This behavior is not desirable so a new definition for the recovery is proposed:

$$\mathrm{I(t)} = \mathrm{I_{ss}} + (\mathrm{I_{peak_1}}-\mathrm{I_{ss}})[1-\exp(-\frac{\mathrm{t}-\mathrm{t_{off_1}}}{\tau_{\mathrm{recov}}})] \tag{4}$$

so

$$\frac{\mathrm{I_{peak_2}}(\tau_{\mathrm{recov}})-\mathrm{I_{ss}}}{\mathrm{I_{peak_1}}-\mathrm{I_{ss}}} = 1-\exp(-1) \tag{5}$$

The opsin currents can be calculated via

$$\mathrm{I(t)} = \mathrm{g_{max}}(\mathrm{O^{on}(t)}+\mathrm{O^{off}(t)})(\mathrm{R^{on}(t)}+\mathrm{R^{off}(t)})(\mathrm{V_m}-\mathrm{E_{ion}}) \tag{6}$$

The following assumptions are made.

$$\mathrm{t_{peak_i}} - \mathrm{t_{on_i}} >> \tau_{\mathrm{O}} \tag{7}$$

$$\mathrm{t_{peak_i}} - \mathrm{t_{on_i}} << \tau_{\mathrm{R}} \tag{8}$$

$$\mathrm{t_{off_i}} - \mathrm{t_{on_i}} >> \tau_{\mathrm{R}} \tag{9}$$

Schoeters et al. (2021) provides an analytic solution for $\mathrm{O^{on}}$, $\mathrm{O^{off}}$, $\mathrm{R^{on}}$ and $\mathrm{R^{off}}$. These can be used in combination with the assumptions to obtain the following approximations:

$$\mathrm{I_{peak,1}} \approx \mathrm{C\,O_\infty} \qquad \text{(assumption (7))} \tag{10}$$

$$\mathrm{I_{peak,2}} \approx \mathrm{C\,O_\infty\,R^{off}(t_{on_2})} \qquad \text{(assumption (7),(8))} \tag{11}$$

$$\mathrm{I_{off_1}} \approx \mathrm{C\,O_\infty\,R_\infty} \qquad \text{(assumption (7),(9))} \tag{12}$$

$$\mathrm{I_{ratio}} \approx \frac{\mathrm{I_{off_1}}}{\mathrm{I_{peak_1}}} = \mathrm{R_\infty} \tag{13}$$

With C = $\mathrm{g_{max}}(\mathrm{V_m}-\mathrm{E_{ion}})$ and

$$\mathrm{R^{off}(t_{on_2})} = 1-(1-\mathrm{R^{on}(t_{off_1})})\exp(-\frac{\mathrm{t_{on_2}}-\mathrm{t_{off_1}}}{\tau_{\mathrm{R}}(\mathrm{I}=0)}) \tag{14}$$

Combining assumptions (8) and (9) implies $\mathrm{t_{peak}}-\mathrm{t_{on}} << \mathrm{t_{off}}-\mathrm{t_{on}}$ so $\tau_{\mathrm{recov}} \approx \mathrm{t_{on_2}}-\mathrm{t_{off_1}}$. The analytic solution of $\mathrm{R^{on}(t_{off_1})}$ in combination with the assumption that steady state is achieved at $\mathrm{t_{off_1}}$ (assumption (9)) results in

$$\mathrm{R^{off}(t_{on_2})} = 1-(1-\mathrm{R_\infty}))\exp(-\frac{\tau_{\mathrm{recov}}}{\tau_{\mathrm{R}}(\mathrm{I}=0)}) \tag{15}$$

Equation (5) can now be written as

$$\frac{1-(1-\mathrm{R_\infty})\exp(-\frac{\tau_{\mathrm{recov}}}{\tau_{\mathrm{R}}(\mathrm{I}=0)})-\mathrm{R_\infty}}{1-\mathrm{R_\infty}} = 1-\exp(-1) \tag{16}$$

$$\Leftrightarrow 1-\exp(-\frac{\tau_{\mathrm{recov}}}{\tau_{\mathrm{R}}(\mathrm{I}=0)}) = (1-\exp(-1)) \tag{17}$$

$$\Leftrightarrow \tau_{\mathrm{R}}(\mathrm{I}=0) \approx \tau_{\mathrm{recov}} \tag{18}$$
